# Mycolactone-independent pathogenicity of *Mycobacterium ulcerans*: an experimental study in plants

**DOI:** 10.1101/867556

**Authors:** A. Bouam, M. Drancourt

**Affiliations:** Aix-Marseille Univ, IRD, MEPHI, IHU Méditerranée Infection, Marseille, France

**Keywords:** Buruli ulcer, *Mycobacterium ulcerans*, tomato, aquatic plants, environment, reservoir

## Abstract

*Mycobacterium ulcerans*, the etiologic agent of Buruli ulcer in humans and animals, secretes macrolide exotoxins mycolactones which damage tissues after a cascade of cellular effects. *M. ulcerans*, an environmental organism with still elusive reservoirs and sources has been detected in soil and water in endemic areas where it could be in contact with plants. Symptom observations, microscopy and molecular biology were used to investigate *M. ulcerans* contact with plants in an experimental model mimicking the known pathology of Buruli ulcer in humans. *Solanum lycopereum* (tomato) plants with scarified or intact roots were transplanted into pots containing contaminated soil with *M. ulcerans* or a mixture of mycolactones A/B and C in the presence of negative control groups. Whereas plants with intact roots remained asymptomatic, *M. ulcerans*-infected plants with scarified roots had significantly more diseased leaves than controls (p = 0.004). Optic microscopy examination showed significantly more mycobacteria in the secondary and main roots than in controls (p=0.0008). Real-time PCRs detected *M. ulcerans* DNA in 7/12 (58%) of infected root samples versus none in the control plants (p = 0.04). Further study of plants with mycolactones A/B and C yielded no significant difference with negative controls. These results suggest that in this model, *M. ulcerans* exhibits a mycolactone-independent pathogenicity whose mechanism remains to be elucidated.

## IMPORTANCE

*Mycobacterium ulcerans* is responsible for a chronic debilitating skin and soft tissue infection that can lead to permanent disfigurement and disability in mammals and humans for whom the so-called Buruli ulcer is the third worldwide most common mycobacterial disease after tuberculosis and leprosy (1). The ultimate environmental reservoir for *M. ulcerans* is unknown but *M. ulcerans* DNA sequences are commonly detected in water and moist soil as well as in biofilms associated to the surface of aquatic and terrestrial plants in endemic tropical countries (2). Furthermore, *M. ulcerans* was shown to survive up to four months in experimentally contaminated moist soil (3) and five environmental *M. ulcerans* isolates have been reported, including one isolate from moss (4–6).

These observations suggest that *M. ulcerans* could naturally be in close contact with plant roots but the relationship between *M. ulcerans* and plant roots has never been studied. Here, using a *Solanum lycopereum* (tomato) model previously reported for *Mycobacterium avium* (7), we explored the interactions of *M. ulcerans* with plant roots and observed that *M. ulcerans* acted as an opportunistic plant pathogen in this model, mimicking what was already known in animals and humans, instating *M. ulcerans* as a trans-kingdom opportunistic pathogen.

## MATERIALS AND METHODS

### *M. ulcerans* strain and mycolactones purification

*M. ulcerans* CU001 was cultured into a Biosafety Level 3 laboratory on Middlebrook 7H10 agar medium supplemented with 10% (v/v) of oleic acid/albumin/dextrose/catalase (OADC) (Becton Dickinson, Sparks, MD, USA) for six weeks at 30°C. IS*2404*, IS*2606* and KR-B based real-time PCR was used to identify the colonies used for the experiments (8). Mycolactones were prepared from *M. ulcerans* CU001 cells as previously described (9,10). Briefly, inactivated colonies were suspended in chloroform-methanol (2:1, v/v). A Folch’s extraction was performed by adding 0.2 volumes of water. The organic phase was collected and dried at 40°C under a stream of nitrogen and phospholipids, which were precipitated by ice-cold acetone. Acetone soluble lipids (ASLs) were then separated by high-performance liquid chromatography (Alliance 2690, Waters, Saint-Quentin-en-Yvelines, France) to collect pure mycolactone A/B and C fractions. ASLs were loaded into a reverse phase column (μBondapak C18 10 μm 3.9×300 mm, Waters), elution was performed at 2 mL/min with 20 % water and 80 % acetonitrile. ASLs were monitored at 363 nm (PDA 996, Waters) and mycolactones A/B and C were individually collected. Mycolactone fractions were further characterized by LC/MS (Acquity iClass UHPLC and Vion IMS Qtof, Waters) and screened for structures against mycolactones A/B, C, D, E, F and G using “Chempsider, UK”. Identified structures were validated using the following criteria: 1) < 5 ppm mass error 2) > 3 predicted fragments 3) < 5 ppm mass error on parent isotopes (root mean square) and 4) < 15 % intensity deviation on parent isotopes (root mean square).

### Experimental infection using viable *M. ulcerans* and mycolactones

A total of 12 adult tomato plants forming four groups of three plants were used for each experiment. The first group was transplanted into pots containing contaminated soil composed of 200 mL of sterilized natural soil mixed with 20 mL of *M. ulcerans* strain CU001 suspension in phosphate-buffered saline (PBS) at 10^5^ colony-forming units CFU/mL final concentration, or a mixture of mycolactones A/B and C. The plants of the second group (negative controls) were transplanted into pots containing sterilized soil mixed with 20 mL of sterile PBS, while the plant roots of the third and fourth group were scarified before transplantation into pots containing contaminated or sterilized soil. Experiments with *M. ulcerans* were conducted in a Biosafety Level 3 laboratory and the transplantations were performed under a laminar airflow hood. Plants were then placed in a hermetic mini greenhouse for the duration of the experiment. After seven days post-transplantation, tomato plants were photographed and the photographs were then coded in order to be read blindly by three different observers who blindly counted the number of diseased leaves. The data were translated into average and standard deviations and the p-value was calculated by comparing groups of contaminated plants with their controls.

### Control of infection

To assess the internalization and the distribution of mycobacteria inside the plants, we chose five sampling points: secondary roots, main root and the lower and the upper part of stems and leaves. The samples were taken aseptically with a sterile scalpel and weighed until they reached 0.5 g, they were then sterilized separately by immersion for three minutes in 30% bleach followed by three rinses in distilled sterile water for 5 minutes, 100 μL of the last rinse was cultured in 7H10-OADC to control the sterility of the sample surfaces. Samples were then crushed using a piston pellet in 2 mL Eppendorf-tubes containing 1mL of sterile PBS and filtered through a 5μm filter to remove the debris. A 1mL-volume of each homogenate was decontaminated with 1% chlorhexidine and then 100 μL were cultured on MOD9 medium (11).

### Optic microscopy

Ziehl Neelsen staining was performed on each homogenate, slides were then coded in order to have blind counts performed by three manipulators and the number of acid fast bacteria (AFB) was counted by analyzing ten fields of observation at a magnification of X1000. The data were translated into average and standard deviations and the p-value was calculated by comparing samples provided from contaminated plants with their controls.

### Molecular biology

Total DNA was extracted manually using Macherey-Nagel extraction kit (Macherey-Nagel, Düren, Germany) following the manufacturer’s instructions. IS*2404* and KR-AB were then tentatively amplified by real-time PCR using RT-PCR reagents (Takyon, Eurogentec, Liège, Belgium) and primers and probes as previously described (8).

### Statistical analyses

Statistical significance was determined using the Student’s t-test. The tests were considered significant if the p-value was inferior to 0.05.

## RESULTS

### *M. ulcerans* provokes a disease in tomato plants

Blind analysis of the plants’ photographs by three independent operators unexpectedly yielded significantly more diseased leaves on contaminated plants with *M. ulcerans* whose roots were scarified compared to their controls in the two experiments (p = 0.004). No significant difference was observed between contaminated plants with non-scarified roots in comparison with their controls (p = 0.43) (Table 1). These results indicated that *M. ulcerans* was provoking a systemic disease in tomato plants which roots had been scarified in this experimental model (Figure 1).

**Table 1:**
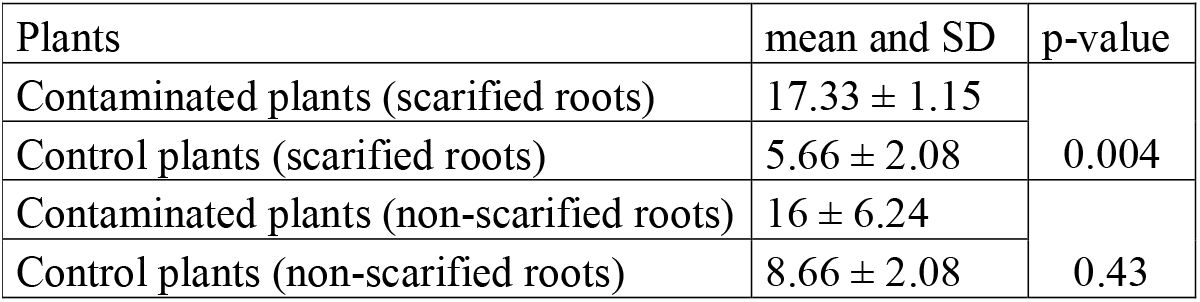
Means and standard deviations of three blind counts of diseased leaves of plants contaminated with *M. ulcerans* and their controls.

| Plants | mean and SD | p-value |
| --- | --- | --- |
| Contaminated plants (scarified roots) | 17.33 ± 1.15 | 0.004 |
| Control plants (scarified roots) | 5.66 ± 2.08 |  |
| Contaminated plants (non-scarified roots) | 16 ± 6.24 | 0.43 |
| Control plants (non-scarified roots) | 8.66 ± 2.08 |  |

**Figure 1.**
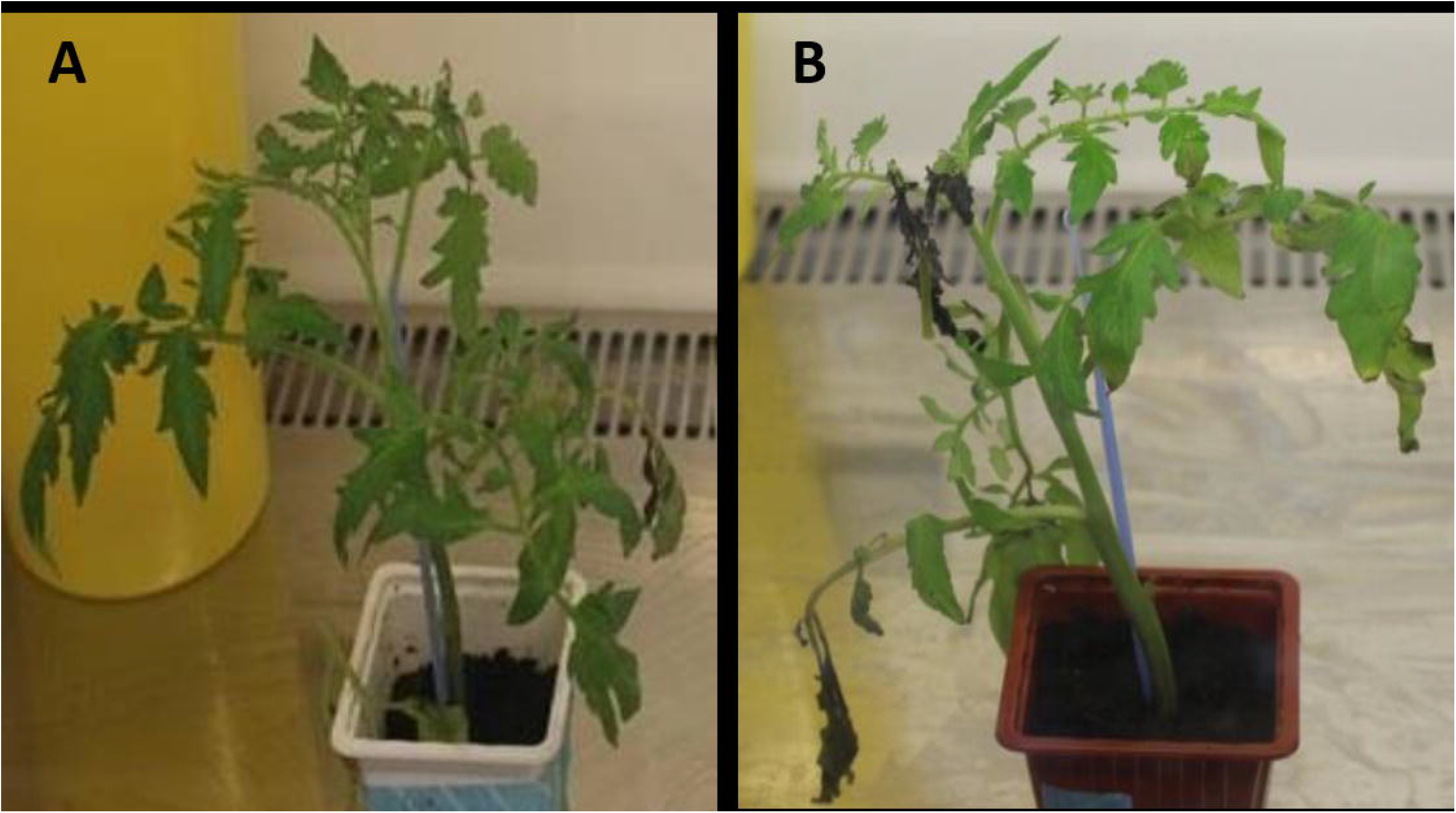
Photography illustrating the effect of *M. ulcerans* on tomato plants. A: control plant. B: *M. ulcerans*-infected plant, exhibiting dark leaves.

### *M. ulcerans* is not invasive in tomato plants

Microscopic examination of secondary roots, the main root and the lower and upper part of stems and leaves specimens after Ziehl-Neelsen staining yielded AFB only in secondary and main roots, but not in the stems and leaves of contaminated plants. No AFBs were observed in specimens collected in negative control plants. The number of AFB observed in the secondary roots of *M. ulcerans*- contaminated plants (99 ± 3) was significantly higher than in negative controls (0 ± 0) (P = 0.0001) (Table 2). Furthermore, the number of AFB observed in scarified secondary roots exposed to *M. ulcerans* (99 ± 3) was significantly higher than in non-scarified secondary roots exposed to the pathogen (70.66 ± 3.51). The number of mycobacteria observed in scarified main roots did not significantly differ from that observed in the non-scarified main roots of pathogen-exposed plants in the duplicate experiment (Table 3). Real-time PCR results confirmed microscopy observations as all stem and leave specimens remained negative in both negative controls and exposed plants. However, *M. ulcerans* DNA was detected in 7/12 (58%; six scarified and one non-scarified) secondary and main roots of *M. ulcerans*-exposed plants and in 0/12 negative controls (p = 0.04, exact Fisher test).

**Table 2:** Means and standard deviations of three blind counts of the number of acid-fast bacilli in secondary and main roots of plants contaminated with *M. ulcerans* and their controls.

| Plants | mean and SD | p-value |
| --- | --- | --- |
| Contaminated plants (scarified secondary roots) | 99 ± 3 | 0.004 |
| Control plants (scarified secondary roots) | 0 |  |
| Contaminated plants (non-scarified secondary roots) | 70.66 ± 3.51 | 0.001 |
| Control plants (non-scarified secondary roots) | 0.33 ± 0.57 |  |
| Contaminated plants (scarified main roots) | 56 ± 6.55 | 0.004 |
| Control plants (scarified main roots) | 0 |  |
| Contaminated plants (non-scarified main roots) | 53.66 ± 3.05 | 0.001 |
| Control plants (non-scarified main roots) | 0.66 |  |

**Table 3:** Means and standard deviations of three blind counts of the number of acid-fast bacilli in secondary and main scarified roots of plants contaminated with *M. ulcerans* in comparison with the number of AFB in secondary and main non-scarified roots of plants contaminated with *M. ulcerans*.

| Plants | mean and SD | p-value |
| --- | --- | --- |
| Contaminated plants (scarified secondary roots) | 99 ± 03 | 0.005 |
| Contaminated plants (non-scarified secondary roots) | 70.66 ± 3.51 |  |
| Contaminated plants (scarified main roots) | 56 ± 6.55 | 0.6 |
| Contaminated plants (non-scarified main roots) | 53.66 ± 3.05 |  |

### *M. ulcerans* disease in plants is independent from mycolactones

After having exposed plant roots to mycolactones A/B and C, blind observations indicated no modification on stems and leaves by comparison with the control group. This observation suggested that *M. ulcerans*-induced systemic disease in tomato plants is not due to any of the tested mycolactones.

## DISCUSSION

The results here reported illustrate that the experimental exposure of tomato plant roots to *M. ulcerans* induces a systemic disease by using a model previously used for *M. avium* experimental infection (7). The results here reported were authenticated by the negative results obtained with negative controls introduced in every step of the experiments and by the concordance of observations made using different methods.

This experimental disease is characterized by yellowing followed by leaf necrosis. The symptoms we observed on the leaves of tomato plants after experimental contamination by *M. ulcerans*. These symptoms were similar to those reported in tomato plants infected by *Burkholderia pseudomallei* (12). Both organisms are environmental microorganisms associated to rice paddies in endemic regions, but in contrast to experimental observations on tomato plants, *M. ulcerans* and *B. pseudomallei* are not yet known to cause diseases in rice plants (12).

The observations here reported indicate that *M. ulcerans* does not provoke an invasive infection, as *M. ulcerans* mycobacteria were associated only with secondary roots and not with any other part of the plant. A previous Illumina technology-based analysis of core actinobacteriome in rice disclosed 7.1 % mycobacteria sequences in roots and a negligible percentage in stems and none in seeds (13). Some bacteria naturally present in the soil, including *Salmonella* spp.*, Escherichia coli* and *Mycobacterium avium*, can colonize plants via their root system (14,15,7). After colonization, some mycobacteria species have benificial or deleterious effects on the plants (16,17).

Our observations suggest that *M. ulcerans*-induced diseased in plants is not due to mycolactones. This observation was unexpected as mycolactones are the only known factor of pathogenicity for *M. ulcerans* (1). It is possible that factors other than mycolactones are implicated in the onset of systemic diseases in experimentally contaminated plants. As for *B. pseudomallei*, plant pathogenicity was linked to the secretion of type three secretion system effectors, a protein secretion in diverse Gram-negative bacteria. Accordingly, we noticed that *M. ulcerans* Agy 99 genome encodes an uncharacterized protein (ABL05176) which contains a region of 78 amino acid named “Secretin”, a bacterial type II and III secretion system protein (18). Alternatively, the symptoms we observed in tomato plants exposed to viable or inactivated *M. ulcerans* organisms may be due to pathogen associated molecular patterns (PAMPS) such as peptidoglycans and lipopolysaccharides, which are known to activate the innate immune response of plants (15). In any case, the tomato plant model here exposed offers a convenient model to further explore the pathogenicity of *M. ulcerans*.

Finally, *M. ulcerans* has to be added to the list of trans-kingdom pathogens. Different studies have shown that, contrary to a previously accepted idea that a strict separation exists between plant and vertebrate pathogens, some plant pathogens may cross the kingdom barrier and be pathogenic for humans such as Pepper mild mottle virus PMMV, reported to cause clinical symptoms and specific immune response in humans (19,20), and the bacterium *Pantoea agglomerans*, responsible for life-threatening catheter-related bacteremia and other conditions in patients (21). Also, human pathogens such as *Salmonella* and *B. pseudomallei* are also reported to cause diseases in plants. In addition, *Salmonella* bacteria use the same mechanism to infect humans and plants and the TSS3s is the virulence factor involved in its pathogenicity for both plants and humans (12,22). Furthermore, some human pathogens such as *Listeria monocytogenes* are tightly associated with plant leaves without causing plant infection (23).

Contrary to these examples of strict pathogens, *M. ulcerans* offers the example of an opportunistic and trans-kingdom pathogen.

## ACKNOWLEDGEMENTS

This work was supported by the French Government under the « Investissements d’avenir » (Investments for the Future) program managed by the Agence Nationale de la Recherche (ANR, fr: National Agency for Research), (reference: Méditerranée Infection 10-IAHU-03). The authors acknowledge the help of Magdalen Lardière and Olga Cusack in English translation and manuscript edition.

